# Dissection of retrosplenial cortex inputs: ubiquitous drive from anterior thalamus

**DOI:** 10.1101/2025.02.06.636939

**Authors:** Gabriella Margetts-Smith, Lilya Andrianova, Shivali Kohli, Andrew D Randall, John P Aggleton, Jonathan Witton, Michael T Craig

## Abstract

The retrosplenial cortex (RSC) is a highly interconnected brain region involved in spatial navigation and associative learning. It forms extensive, reciprocal connections with sensory, hippocampal, parahippocampal, prefrontal, and thalamic areas. RSC comprises granular (gRSC) and dysgranular (dRSC) subdivisions with distinct connectivity and functions. Despite its emerging role in behaviour and its implication in memory-related disorders such as Alzheimer’s disease, the nature of its synaptic inputs remains poorly understood.

Here, we combined viral anatomical tracing, optogenetic stimulation, and patch-clamp electrophysiology to investigate inputs from the anterior cingulate cortex (ACC), dorsal subiculum (dSub), and anterior thalamic nuclei (ATN) to gRSC and dRSC. Strikingly, all recorded RSC pyramidal neurons received ATN input, regardless of subdivision or cortical layer. Activation of ATN inputs evoked significantly larger post-synaptic responses than those from dSub or ACC, though both regions maintained substantial connectivity with RSC. While dSub projections appeared denser in gRSC, synaptic responses were larger in dRSC, albeit with lower input probability. Notably, NMDA receptor-mediated components of RSC excitatory inputs were weaker than expected, potentially explaining the reported inability to induce long-term potentiation in RSC in *ex vivo* neurophysiology experiments.

This is the first study to characterise the synaptic properties of retrosplenial afferents. Our findings highlight the dominant influence of ATN inputs and raise important questions about how RSC’s long-range connectivity supports its roles in memory and spatial navigation.

## Introduction

Spatial navigation and memory are cognitive functions that have long been associated with the hippocampal formation. Research in recent decades has, however, revealed that several other brain regions, including the anterior thalamic nuclei (ATN) and retrosplenial cortex (RSC), play key roles in these processes (reviewed by [1]). The RSC is extensively and reciprocally connected with various sensory cortices and thalamic nuclei, and also exhibits connectivity with hippocampal and parahippocampal structures. This pattern of connectivity is conserved from mouse to human. This suite of connectivity likely underlies the important integrative function of the RSC enabling the formation of the complex representations necessary for navigating the world; such as the integration of allocentric and egocentric spatial information [2,3].

The rodent RSC is typically divided into 2 subdivisions, based on the presence or otherwise of layer 4; the more ventral subdivision is called granular retrosplenial cortex (gRSC) while the dorsal subdivision is referred to as dysgranular retrosplenial cortex (dRSC) [2]. Functionally, the ATN and hippocampus provide key inputs to the RSC. Removal of ATN input via experimental lesion results in a loss of the ability of RSC to express synaptic plasticity [4], whereas loss of hippocampal input drives widespread reductions in c-Fos expression in RSC [5]. Furthermore, suppression of subicular input impairs flexible spatial memory [6]. This connectivity likely has functional consequences in humans: RSC is one of the first brain regions to show reduced metabolism in Alzheimer’s disease [7,8].

RSC is one of the largest regions in the rodent brain, emphasising its diverse and important functions. Much of what we know about RSC connectivity with the rest of the rodent brain comes from neuroanatomical studies using classical tracers in rats. Such studies reveal differences in connectivity between gRSC and dRSC, but also highlight connectivity with ATN as a common feature to both subdivisions [9–12]. Hippocampal inputs to RSC, arising from the dorsal subiculum, send a denser projection to gRSC than dRSC, and at least half of subicular neurons projecting to gRSC also send collaterals to the mamillary bodies [13]. It should be noted that there are also widespread reciprocal connections within RSC itself [14]. However, as we previously found when studying connectivity of prefrontal cortex and thalamic nucleus reuniens with hippocampus [15,16], relying on anatomical studies of projections alone is not sufficient to understand long-range connections between different brain regions. A combination of optogenetics and patch-clamp electrophysiological recordings can provide novel insights into functional connectivity at the synaptic and cellular level.

Glutamatergic neurons are the principal drivers of excitation throughout the cortex and hippocampus. Electrophysiological studies have revealed substantial heterogeneity in excitatory pyramidal cells (PCs) within RSC. Within superficial layers (layer 2 & 3) of gRSC, small PCs with a late-spiking phenotype have been reported in rat [17] and, more recently, in mice [18]. Larger, burst-spiking PCs can be found in layer 5 of rat gRSC [19]. Recent *in vivo* studies have provided significant advances in our understanding of the functional properties of RSC neurons, particularly dRSC, during navigational and cognitive behaviours (e.g. [20]), but the synaptic inputs that drive cellular responses remain poorly categorised. ATN projections to gRSC have been shown to target the apical dendrites of layer 5 PCs, where they also interact with inhibitory projections from dorsal CA1 [21]. Another recent mouse experiment, using current-clamp recordings, found that the ATN target both late- and regular spiking PCs in gRSC, and that both anterior cingulate and claustrum inputs provided stronger drive to regular-spiking relative to late-spiking PCs, while dorsal subiculum inputs displayed the converse [22].

Little is known, however, of the synaptic properties (typically measured in voltage clamp recordings) of different afferent inputs to gRSC, and we are not aware of any published description of inputs to dRSC despite significant recent advances in knowledge of the functional profiles of neurons in this area during animal behaviour. Here, using a combination of optogenetic circuit-mapping and voltage clamp electrophysiological recordings, we sought to identify and compare the specificity and synaptic properties of inputs to RSC from ATN, dorsal subiculum (dSub) and the anterior cingulate cortex (ACC), for both dRSC and gRSC.

## Results

### Anatomical characterisation of different afferent inputs to the RSC

We used anterograde viral tracing with eGFP driven by the *hSyn* promoter to examine differences in afferent targeting of the mouse RSC from three different upstream regions: the anterior cingulate cortex (ACC; Supplementary Figure 1A), the dorsal subiculum (dSub; Supplementary Figure 1B) and anterodorsal and anteroventral thalamic nuclei (AD/AV; Supplementary Figure 1C). Fluorescence from dSub and AD/AV projections was only seen in the ipsilateral RSC; ACC input showed preferential projection to the ipsilateral RSC, with negligible fluorescence observed in the contralateral RSC (Figure 1A & Supplementary figure 2). Therefore, all further analyses describe afferent terminal fluorescence in the ipsilateral RSC only. Furthermore, no significant difference in afferent terminal distribution was observed along the longitudinal axis of the RSC, so data from anterior and posterior sections has been combined.

**Figure 1.**
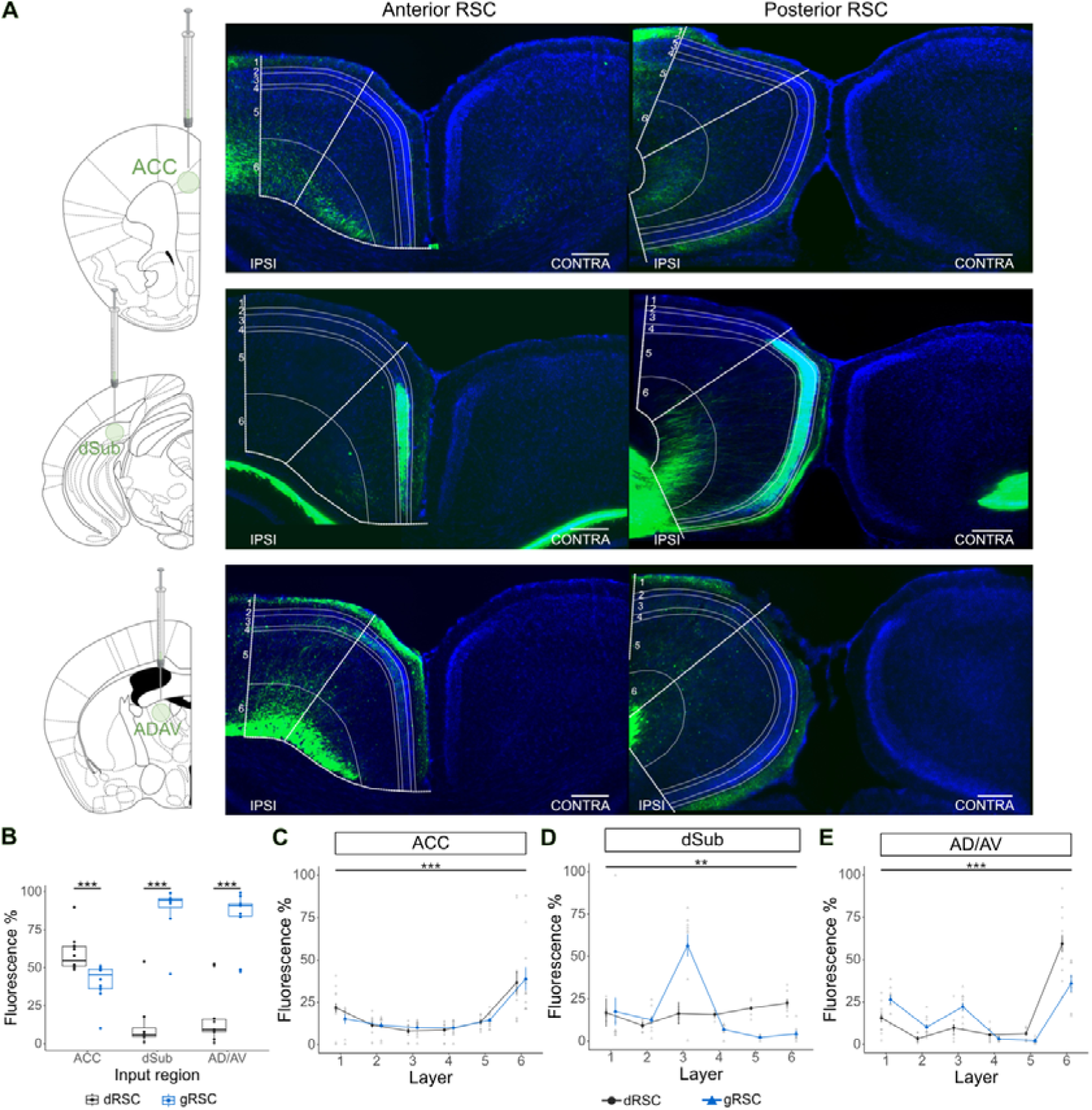
Anterograde viral tracing from the ACC, dSub and AD/AV shows distinct afferent terminal distributions in the RSC. **A,** representative images of EGFP-labelled afferent terminals in the anterior (left) and posterior (right) RSC for each presynaptic region. Scale bars: 250 µm. **B,** there was a main effect of RSC sub-region (*H* (1) = 11.02, *p* < .001; Scheirer-Ray-Hare test), and a significant interaction between sub-region and presynaptic region (*H* (2) = 29.73, *p* < .001). Mann-Whitney U *post hoc* pairwise comparisons revealed significantly higher fluorescent signal in the gRSC compared to the dRSC for inputs from the dSub and AD/AV, whilst ACC input showed higher fluorescent signal in the dRSC compared to the gRSC (all *p* < .001). **C,** there was a significant main effect of layer for ACC input (*F* (5,110) = 19.5, *p* < .001; 2-way mixed ANOVA with a Greenhouse-Geisser correction), but no interaction between sub-region and layer (*F* (5,110) = 0.45, *p* = .53). **D**, there was a significant main effect of layer for dSub input (*F* (5,70) = 8.30, *p* < .01; 2-way mixed ANOVA with a Greenhouse-Geisser correction) and a significant interaction between sub-region and layer (*F* (5,70) = 9.69, *p* < .01). **E,** there was a significant main effect of layer for ATN input (*F* (5,90) = 64.08, *p* < .001; 2-way mixed ANOVA with a Greenhouse-Geisser correction) and a significant interaction between sub-region and layer (*F* (5,90) = 10.41, *p* < .01). Boxplots display median, IQR and range. Line-graphs display mean ±SEM. *\* p < .05, ** p < .01, *** p < .001*.

We found significant differences in afferent terminal distribution between the dRSC and gRSC for all presynaptic regions (Figure 1B). Inputs from the ACC preferentially targeted the dRSC, whereas inputs from the dSub and AD/AV preferentially targeted the gRSC. Subsequent laminar analyses of fluorescence intensity (see methods) indicated that ACC axonal innvervation was most dense in layer 6 in both the dRSC and gRSC (Figure 1C). Conversely, afferents from both the dSub and AD/AV preferentially targeted different cortical layers between RSC sub-regions. Terminals from the dSub were largely uniformly distributed in the dRSC, but there was strong preference for layer 3 in the gRSC (Figure 1D). AD/AV terminals were predominantly observed in the dRSC layer 6, with a smaller percentage of fibres also present in layers 1 and 3 of the gRSC (Figure 1E).

### Functional synaptic input probability and amplitude differs between RSC afferent pathways

We observed similar sub-region and cortical layer targeting in the mouse RSC compared to previous studies of the rat brain [9–11], with some key differences such as subicular input into the dRSC in the murine brain. However, confirming structural connectivity between these input regions and the RSC does not necessarily confer insight into their functional connectivity, as we found for nucleus reuniens [15]. Therefore, we next measured synaptic connectivity of ACC, dSub and AD/AV afferents onto pyramidal cells (PCs) in the dRSC and gRSC. To measure the synaptic connectivity of major afferent pathways in RSC, we injected a viral vector to drive expression of channelrhodpsin2 (ChR2) in ACC, AD/AV or dSub, respectively, and subsequently stimulated the afferent terminals while conducting *ex vivo* voltage clamp whole-cell recordings in RSC PCs (Figure 2A).

**Figure 2.**
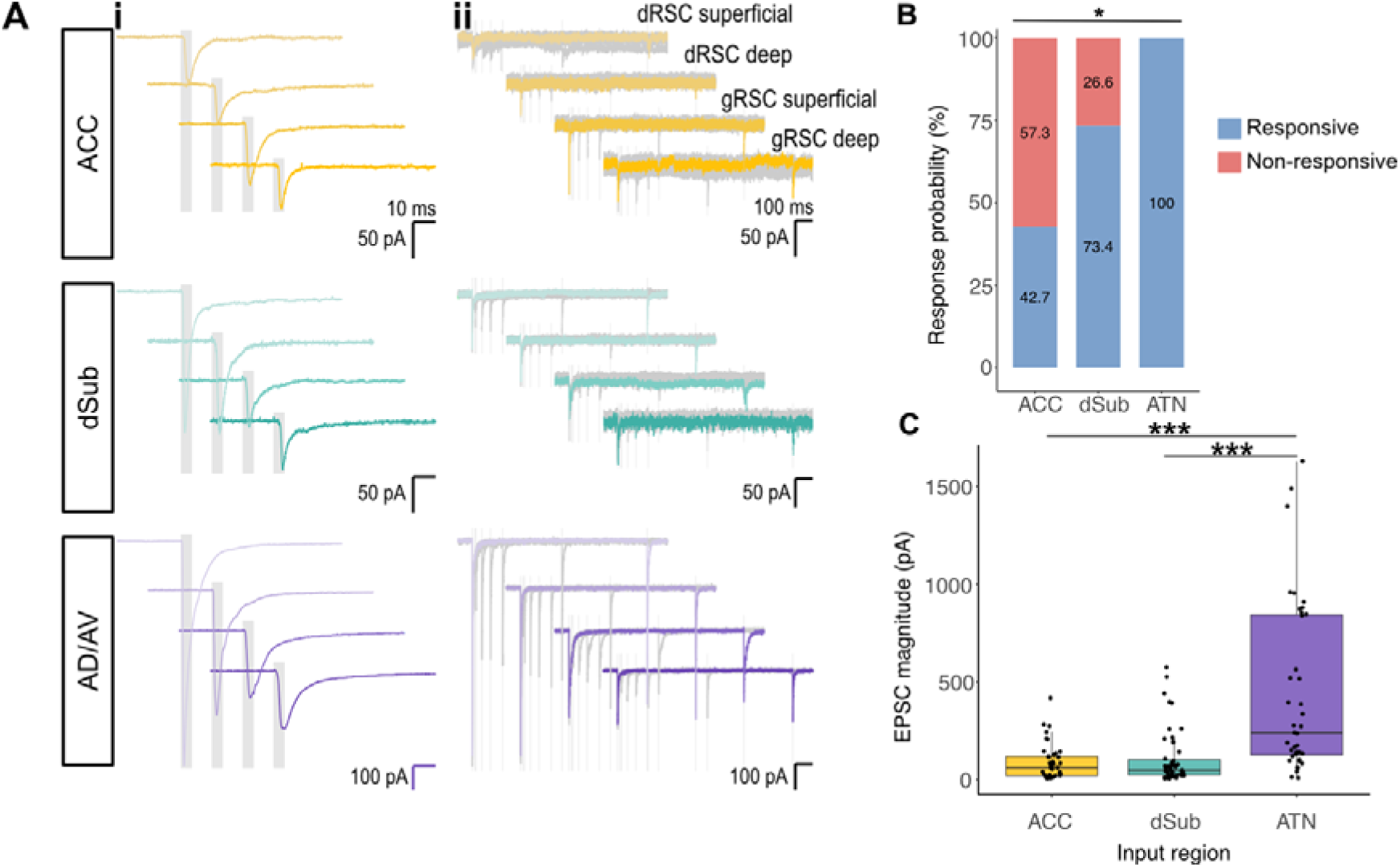
Synaptic response in the RSC differs between input regions. **A**, Representative voltage clamp (V_H_ = −70 mV) traces. (**i**) EPSCs generated in the first optogenetic stimulation of a 5 pulse 7 Hz stimulation protocol. (**ii**) EPSCs generated by two optogenetic stimulations separated by increasing intervals. Each plot displays 7 sweeps with intervals of (grey traces) 10 ms, 17 ms, 51 ms, 100 ms, 170 ms, 510 ms and (coloured trace) 1000 ms. Optical stimulation periods indicated by grey boxes. **B**, probability of a synaptic response in RSC pyramidal cells significantly differed dependent on presynaptic region (χ^2^(2) = 43.20, *p* < .001; Pearson’s Chi-squared test). Chi-squared *post hoc* tests showed each presynaptic region elicited a different response probability (all *p* < .001) with ACC input generating the lowest probability, and AD/AV input the highest. **C**, there was a main effect of presynaptic region on median EPSC magnitude (*H* (2) = 33.28, *p* < .001; Kruskal-Wallis test). Mann-Whitney U *post hoc* tests showed EPSCs generated from ACC and dSub input did not differ, EPSCs generated from ATN were significantly larger (*p* < .001). *\* p < .05, *** p < .001*.

Considering RSC in its entirety, we found that while all tested inputs generated an excitatory postsynaptic current (EPSC) in RSC pyramidal cells, there was a significant difference in probability of response. Input from the ACC elicited the lowest response probability (42.7%), followed by input from the dSub (73.4%), and finally AD/AV input elicited an EPSC in 100% of the PCs recorded (Figure 2B). In addition to differing probabilities of response, we found the different presynaptic inputs elicited different amplitudes of excitatory AMPA receptor-mediated currents (Figure 2C). Averaged across the RSC, inputs from the AD/AV generated a significantly larger ESPC (∼7-9 fold) than inputs from the ACC and dSub, which did not differ in amplitude. We next conducted mixed model analyses to investigate the effect of PC location on elicited EPSC amplitude. Interestingly our results indicate a disparity between the structural connectivity demonstrated in Figure 1B and strength of excitation for each tested input.

We parsed RSC PCs into superficial (supragranular) and deep (infragranular) in gRSC or dRSC, respectively, to take account of RSC having two major subdivisions containing heterogeneous PC subtypes, as classified elsewhere by firing properties (e.g. [17–19]). We found that the probability of response to afferent excitation also differed between sub-region and layer for ACC inputs, but not for dSub inputs (Figure 3A – B). The majority of superficial neurons in gRSC received no input from ACC, while there appeared to be no difference in selectivity for deep *vs* superficial in dRSC. As all recorded RSC neurons received input from AD/AV, we did not further subdivide these data for AD/AV input probability.

**Figure 3:**
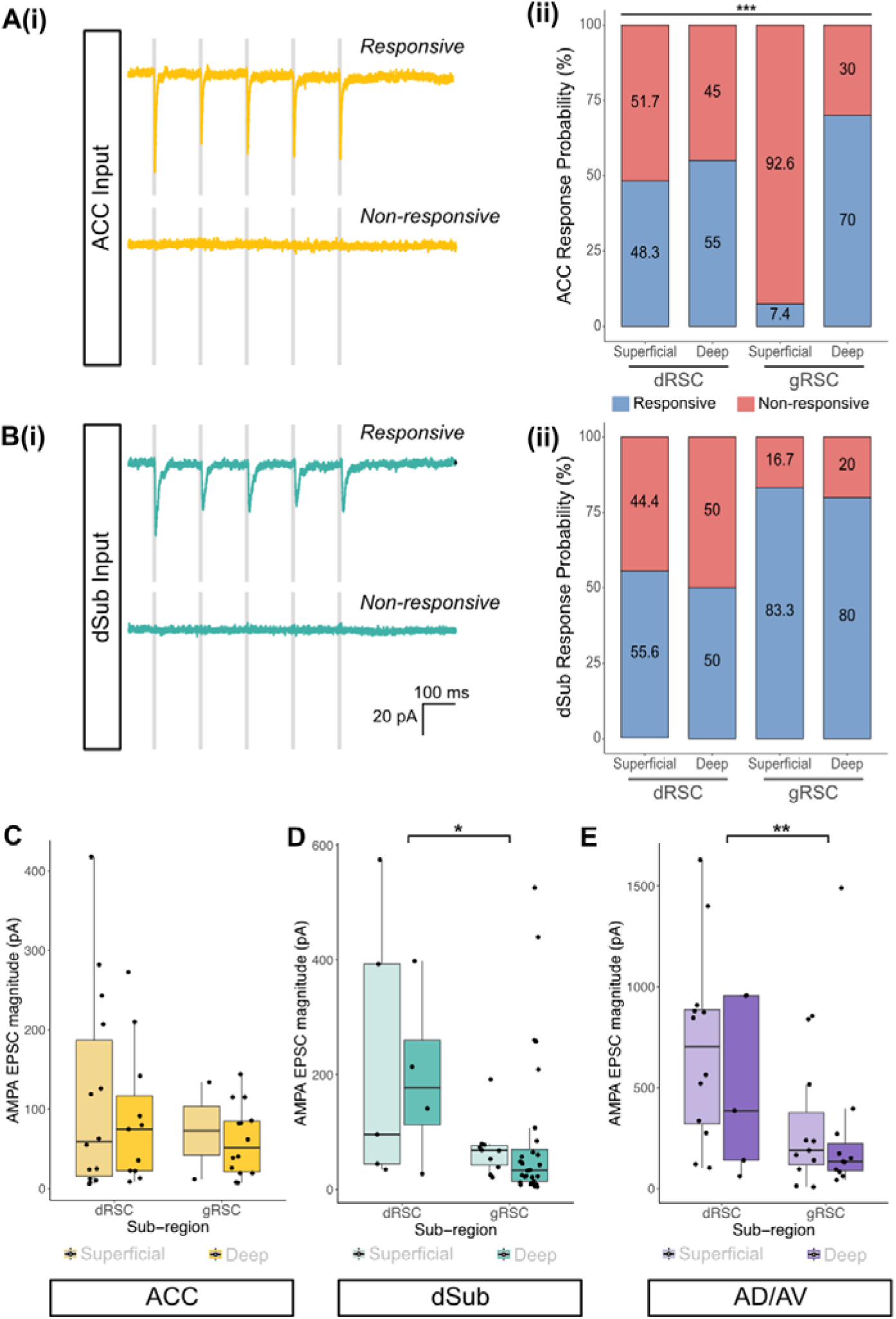
Afferent inputs to RSC, broken down by subdivision and laminar location. **A**, probability of EPSC response following ACC axon terminal stimulation in different sub-regions and cortical layers of the RSC. (**i**) Representative voltage-clamp (V_H_ = −70 mV) traces of responsive and non-responsive PCs recorded in the superficial dRSC. (**ii**) Probability of PC synaptic response following ACC input stimulation differed significantly dependent on cell location within the RSC (χ*^2^* (3) = 21.44, *p* < .001; Pearson’s Chi-squared test). Chi-squared *post hoc* tests revealed that superficial gRSC cells had a significantly lower response probability than those in the deep gRSC (*p* < .001) and both superficial and deep dRSC cells (both *p* < .01). The other groupings did not significantly differ from one another. **B**, probability of EPSC response following dSub axon terminal stimulation in different sub-regions and cortical layers of the RSC. (**i**) Representative voltage-clamp (V_H_ = −70 mV) traces of responsive and non-responsive PCs recorded in the superficial dRSC. (**ii**) PC location within the RSC did not significantly affect probability of EPSC response following stimulation of dSub axon terminals (χ*^2^* (3) = 5.10, *p* = 0.16; Pearson’s Chi-squared test). **C**, for ACC input, RSC sub-region or layer did not significantly affect EPSC magnitude. **D**, for dSub input, EPSC magnitude was significantly higher in cells in the dRSC than in the gRSC, but there was no difference between cells in superficial vs deep cortical layers. **E**, for ADAV input, EPSC magnitude was significantly higher in cells in the dRSC than in the gRSC. But there was no difference between cells in the deep vs superficial cortical layers. *\*\*\* p < .001*.

Despite the anatomical tracing suggesting that ACC preferentially innervates dRSC, there was no significant difference in AMPA receptor-mediated EPSC amplitude between dRSC and gRSC, or between superficial and deep cortical layers within each sub-region (Figure 3C). Furthermore, whilst dSub and AD/AV afferent terminals were markedly denser in the gRSC, both inputs elicited greater excitatory currents in the dRSC (Figure 3D & 3E) compared with gRSC. Full statistical results of the mixed effect model of EPSC magnitude for each input are presented in Supplementary Table 1. Within both dRSC and gRSC, the magnitude of AMPA receptor-driven afferent synaptic inputs was comparable for deep and superficial PCs (Figure 3C – E).

### Indicators of synaptic plasticity are consistent between RSC afferent pathways, but differ between RSC sub-region and cortical layer

Next, we examined the NMDA receptor-mediated component of the EPSC from each afferent input to RSC (Figure 4A), as NMDA receptors are a key mediator of long-term synaptic plasticity (e.g. [23]). We found that the ACC input evoked the smallest NDMAR- EPSC, with that from dSub being significantly larger; and - as might be expected given its larger AMPAR-mediated component (Figure 2C) - the AD/AV input evoked a significantly larger NMDAR-EPSC than both ACC and dSub inputs (Figure 4B). We then compared the ratio of AMPAR- to NDMAR-evoked EPSCs, and observed a low NMDA/AMPA receptor ratio for all inputs (Figure 4C), which did not differ significantly between these afferent pathways, RSC subdivision, or laminar location (Figure 4D – F). Our observation of low NMDAR/AMPAR ratios elicited from all inputs support previous reports that LTP cannot be induced in the RSC following tetanic stimulation [4]. Full statistical results of the NMDAR/AMPAR ratio mixed models are presented in Supplementary Table 3.

**Figure 4:**
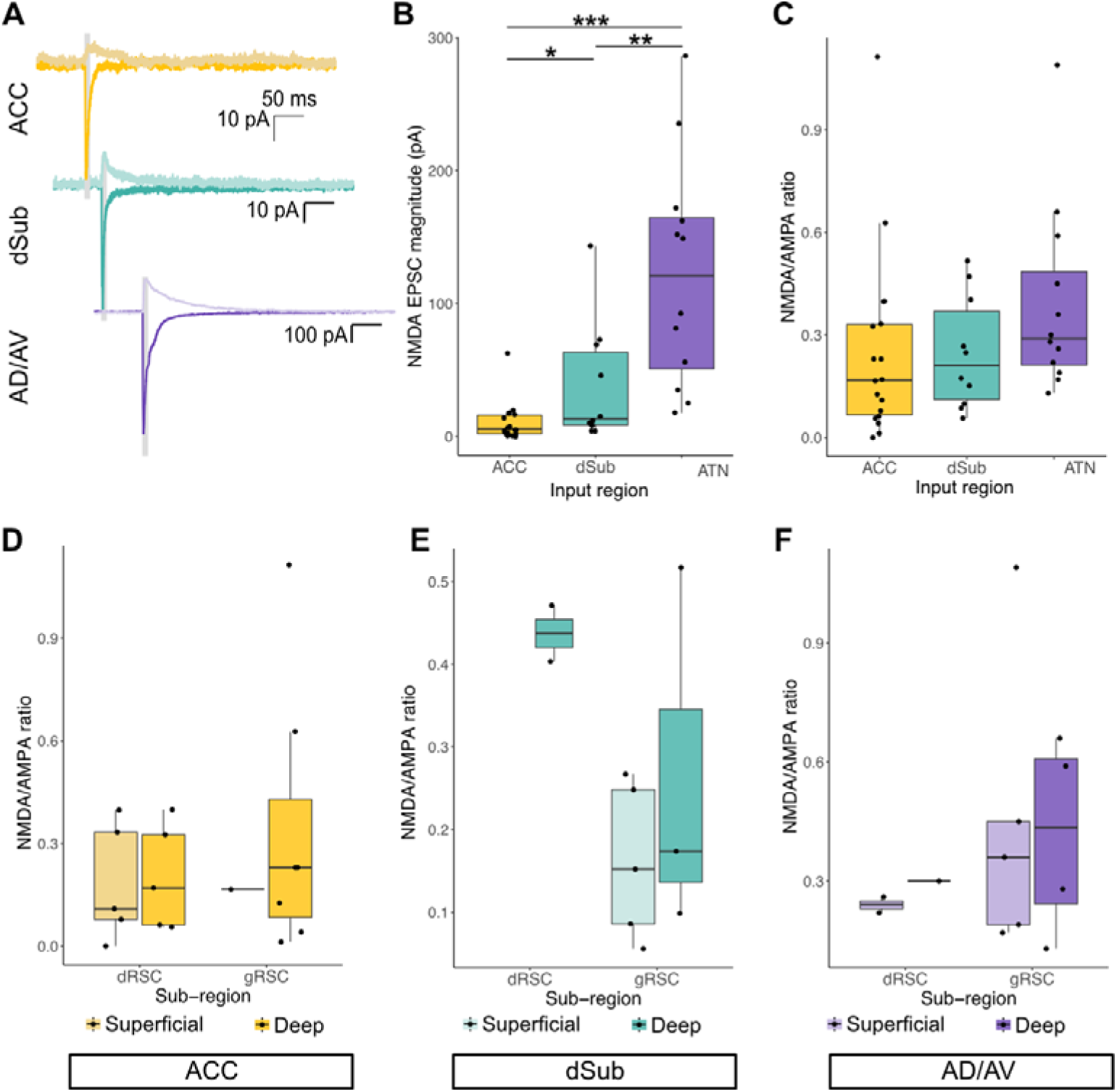
NMDA-receptor mediated currents on afferent inputs to RSC. **A,** representative voltage-clamp traces showing AMPAR-mediated (dark) and NMDAR- mediated (light) EPSCs. AMPAR responses were recorded at V_H_ = −70 mV (standard aCSF) and NMDAR responses were recorded at V_H_ = +40 mV (standard aCSF containing (in µM): 10 DNQX, 1 CGP-5584, 10 Gabazine). **B**, NMDA-mediated EPSC magnitude significantly differed between presynaptic regions (*H* (2) = 21.78, *p* < .001). **C**, NMDA/AMPA ratio did not differ significantly between presynaptic regions (*H* (2) = 4.28, *p* = 0.12; Kruskal-Wallis test). **D – F**, for all inputs, RSC sub-region or layer did not significantly affect NMDA/AMPA ratio. * *p < .05, ** p < .01, *** p < .001*.

Finally, we examined short-term plasticity by determining paired pulse ratio (PPR), using a protocol of increasing inter-pulse-intervals (IPI). Facilitation or depression of the second EPSC amplitude is indicative of distinct presynaptic dynamics related to short-term plasticity [24], and our results showed significant paired-pulse depression (mean PPR < 1) following stimulation of all three presynaptic inputs (Figure 5A – B). PPR did not vary significantly by either laminar location or RSC subdivision for ACC inputs (Figure 5C). However, paired pulse ratio was higher in the gRSC and deep cortical layers for both dSub and AD/AV inputs (Figure 5D – E). Interestingly, the amplitude of the EPSC did not appear to correlate with subsequent depression as EPSC amplitudes were higher in the dRSC than gRSC for both pathways. Supplementary Table 2 details complete statistical results for each mixed model effect model of PPR. These findings suggest that whilst these measures of synaptic plasticity do not differ overall between afferent inputs to the RSC from the ACC, dSub and AD/AV, the strength of short-term paired pulse depression is dependent on postsynaptic soma location for dSub and AD/AV, but not ACC, inputs.

**Figure 5:**
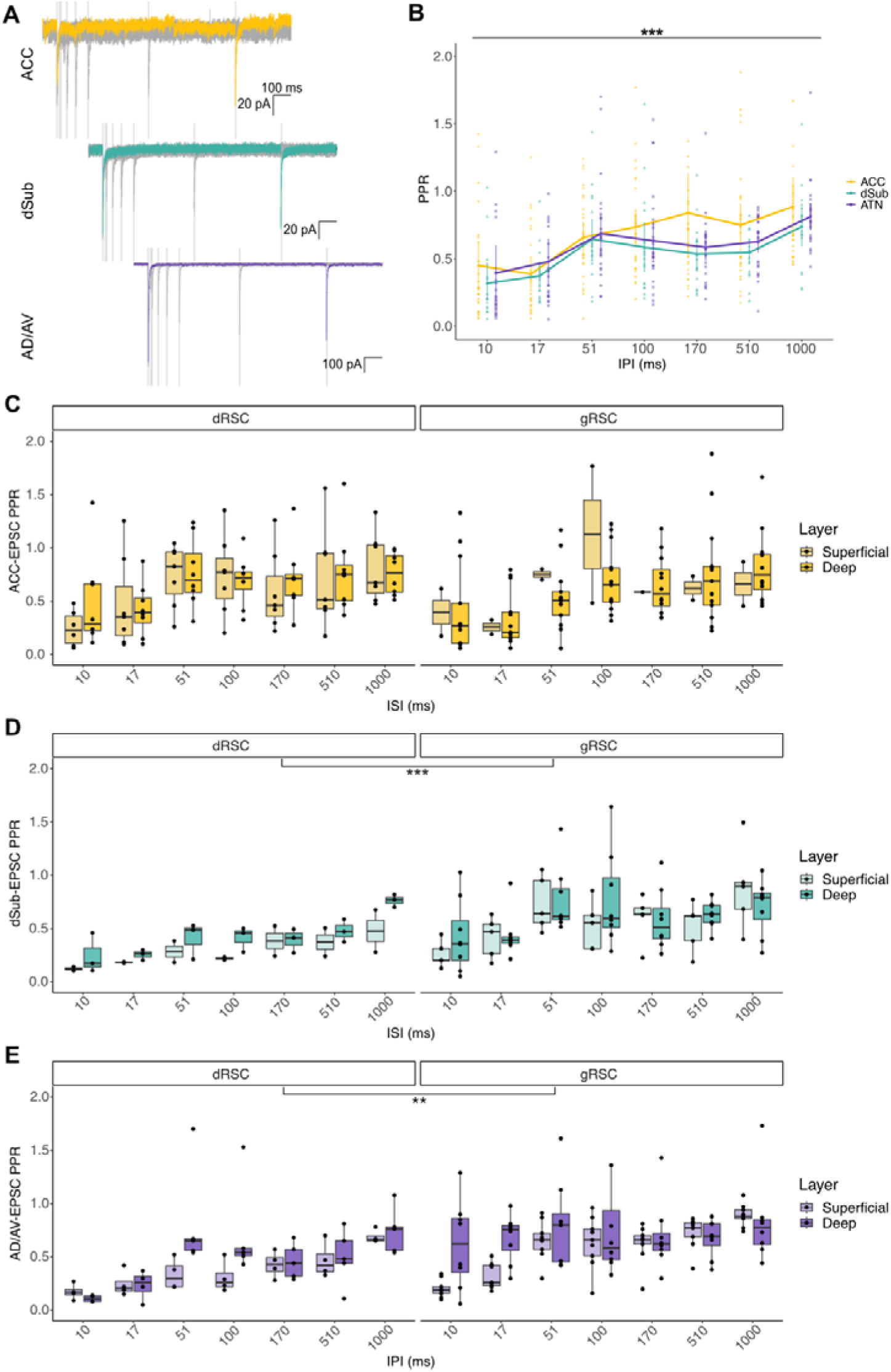
RSC afferent input paired pulse ratio (PPR). **A**, representative voltage-clamp (V_H_ = −70 mV) traces showing EPSCs generated by two stimulations separated by increasing intervals. Each trace displays 7 sweeps with intervals of (dark grey) 10 ms, 17 ms, 51 ms, 100 ms, 171 ms, 510 ms and (black) 1000 ms. 5 ms optical stimulation periods indicated by light grey boxes. **B**, there was a significant main effect of IPI on PPR (*F* (6,426) = 15.71, *p* < .001; 2-way mixed ANOVA), with a steady increase in ratio as IPI increased. There was, however, no overall significant difference between the presynaptic regions (*F* (2, 71) = 1.67, *p* = .22), nor an interaction between presynaptic region and IPI (*F* (12, 426) = 1.02, *p* = .43). Line graph displays mean ±SEM. **C**, for ACC input, PPR increased as IPI increased, but there was no significant effect of sub-region or cortical layer on PPR. **D**, for dSub input, overall PPR increased as IPI increased. Both sub-region and layer significantly affected PPR, with higher PPR seen in gRSC vs dRSC cells, and deep vs superficial cortical layers. **E**, for AD/AV input, overall PPR increased as IPI increased. Both sub-region and layer significantly altered PPR, with higher PPR seen in gRSC vs dRSC cells, and deep vs superficial cortical layers. Boxplots display median, IQR and range. Black dashed line indicates PPR = 1. *\*\* p < .01, *** p < .001*.

### Heterogeneity of pyramidal cells (PCs) in the RSC

The findings above suggest that functional connectivity into the RSC differs not only between origin of input but is also dependent on PC location within the RSC. Prior studies have identified substantial heterogeneity of PC subpopulations within the gRSC [17–19] and layer 5 dRSC [25], yet there is still relatively little known about PC subpopulations in the RSC compared to many other brain regions. We therefore characterised and compared the active and passive intrinsic properties of superficial and deep layer PCs across both the dRSC and gRSC. We expected that the membrane and firing properties of PCs in the RSC would also differ dependent on laminar and sub-region locality, alongside the differences in the macro-circuitry described above.

We analysed 12 intrinsic physiological properties (Table 1 and Figure 6A) using a hierarchical clustering model, from which 3 discrete clusters emerged (Figure 6B). Clusters 1-3 (C1-3) appeared relatively distinct, but when using a PCA, we observed an overlap between C1 and C3 (Supplementary Figure 3A). Additionally, C2 and C3 were comparatively quite dispersed, with a low Dunn Index value of 0.11, indicating a probable high intra-cluster variability and potential for further sub-typing within these clusters in future. Bootstrapping indicated high stability for C1 (> 0.85) and moderate stability for C2 and C3 (> 0.7) (C1: stability = 0.90, dissolutions = 2; C2: stability = 0.65, dissolutions = 42; C3: stability = 0.76, dissolutions = 7). While we found that the variance explained by 3 clusters was fairly low (k_3_ = 30.0%), it was a substantial increase from 2 clusters (k_2_ = 18.5%). Furthermore, although variance did continue to increase in smaller increments for higher cluster numbers (k_4_ = 35.1%, k_5_ = 39.6%), plotting via PCA indicated k_>3_ were highly overlapping and indistinct from each other (data not shown). Example spiking properties and *post hoc* morphological recoveries of neurons from different clusters are shown in Figure 6D – I.

**Figure 6:**
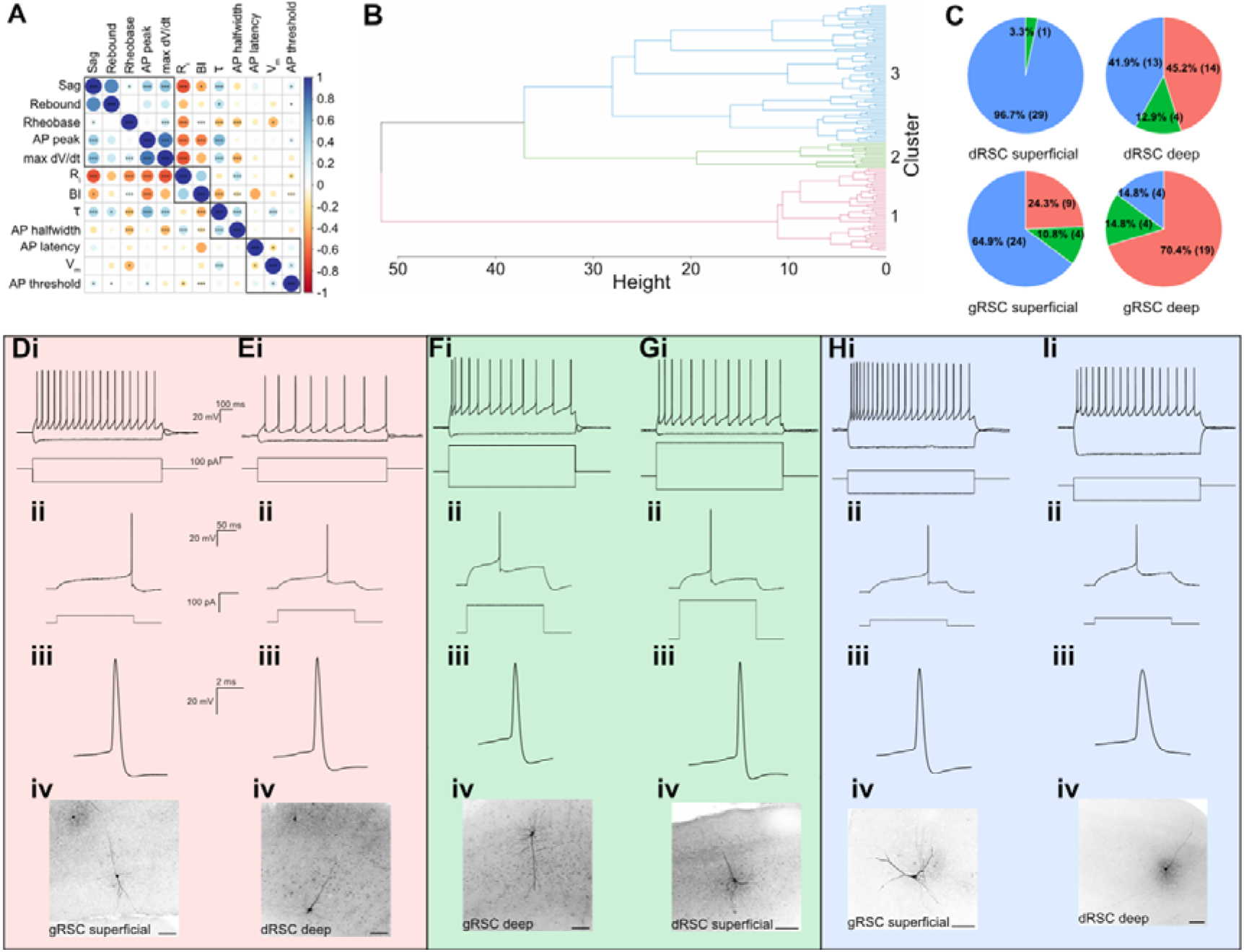
PCs in the RSC are highly heterogeneous. **A**, there was a large amount of correlation between the non-normalised variables, significance values indicated on correlogram. **B**, Dendrogram showing the hierarchical clustering of PCs recorded in the RSC. **C**, RSC sub-regions contain different ratios of each cell cluster type. PCs in the dRSC superficial layer region showed a significant difference in cluster allocation (χ*^2^* (2) = 54.20, *p* < .001); no cells belonged to cluster 1, while the majority belonged to cluster 3. However, ratios of PCs in each cluster group did not significantly differ in the dRSC deep layer region (χ^2^ (2) = 5.88, *p* = .05). PCs in the gRSC superficial layers showed a significant difference in cluster allocation (χ*^2^* (2) = 17.57, *p* < .001); the largest percentage of cells belonged to cluster 3. Finally, PCs in the gRSC also showed a significantly different cluster allocation ratio (χ^2^ (2) = 16.67, *p* < .001), with the highest frequency of cells belonging to cluster 1. Pie charts are annotated with cell percentage and count (in brackets) for each cluster. **D – I**, example current clamp traces for neurons in C1 (**D & E**), C2 (**F & G**) and C3 (**H & I**), showing (**i**) voltage response (top) in response to depolarising current injections sufficient to induce >4 spikes; (**ii**) action potential firing at rheobase; **iii)** first spike in train in i on an expanded time base; (**iv**) post hoc recovery of the neuron morphology using staining for biocytin. Scale bar: 100 µm.

**Table 1.**
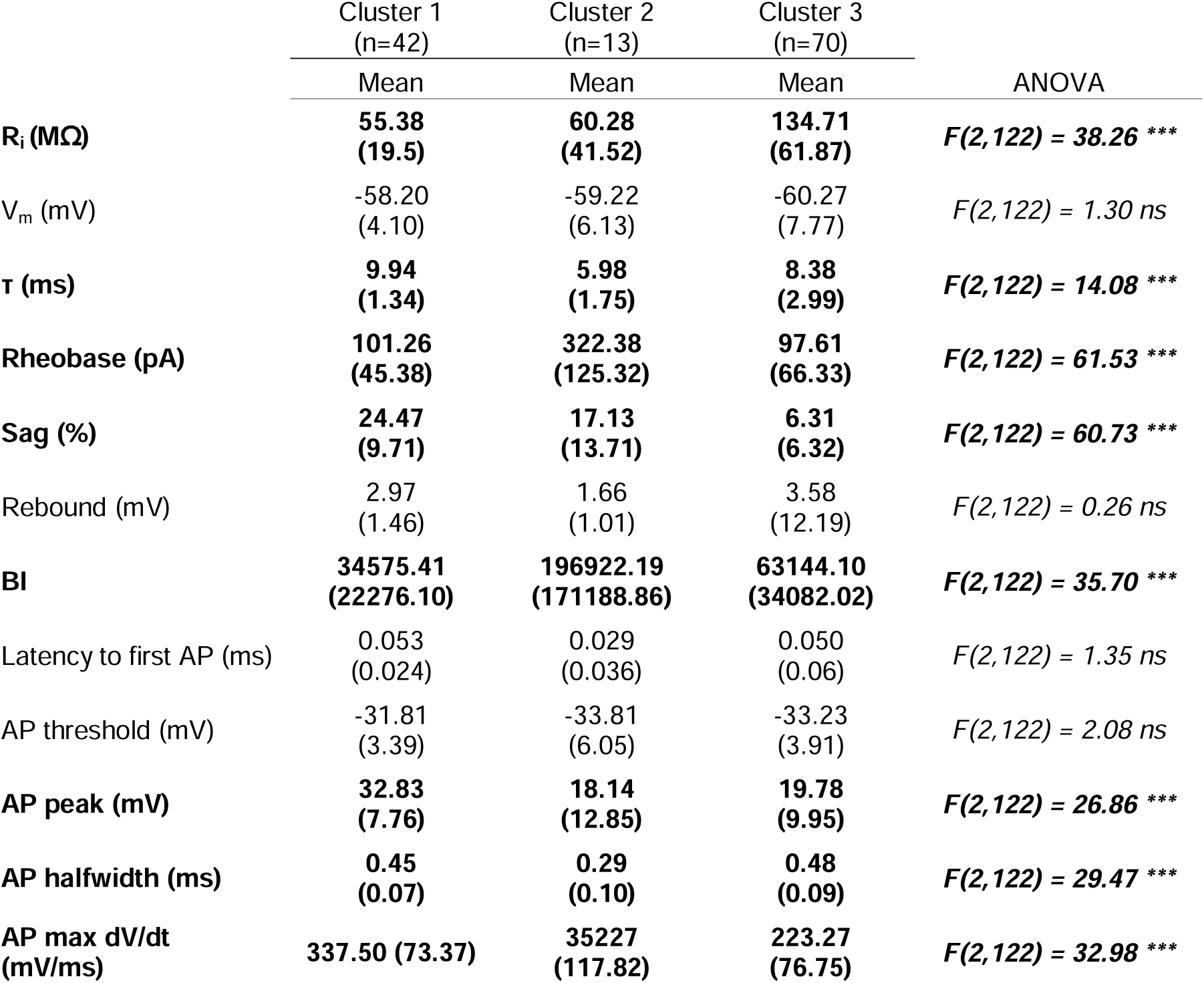
Summary of intrinsic membrane properties between clusters. Mean and SD deviation are shown for each cluster, and univariate ANOVA results are presented. Significant differences between clusters were found for R_i_, rheobase, sag, BI, AP peak, AP halfwidth and AP max dV/dt. *** *p < .001*.

Consideration of clustering upon principal component 1 and 2 axes (Supplementary Figure 3A) and associated variable loadings (Supplementary Figure 3B) suggested that input resistance (R_i)_, membrane time constant (τ), rheobase, burst index (BI), action potential (AP) max dV/dt and actional AP peak amplitude had the strongest influence on cluster allocation. This was supported by the significant differences we observed between clusters in these measures (Supplementary figure 3D). As the non-standardised variables showed a high collinearity, we conducted an initial MANOVA which showed that intrinsic physiological properties differed significantly between the three clusters (Pillai’s trace = 1.38, *F* (24,224) = 20.64, *p* < .001). We followed this up with univariate ANOVA (Table 1) - and additional *post hoc* tests - to identify where these differences appeared within the results. Overall, we found that cells in C1 presented with the highest τ, sag and AP peak as well as the lowest BI. C2 cells had highest rheobase and the lowest τ and AP halfwidth, and C3 cells the highest R_i_ and the lowest sag and AP max dV/dt (Supplementary Figure 3D).

Thus, our results reveal three distinct clusters of RSC PCs, identifiable by specific membrane and firing properties. We next explored the distribution of these putative PC subtypes within the RSC (Figure 3C) and found significantly different ratios of each cell cluster between the dRSC and gRSC (χ*_2_* (6) = 43.92, *p* < .001; Pearson’s Chi-squared test). *Post hoc* tests revealed that PC subtype distribution further differed dependent on cortical layer as well as RSC subregion (Figure 3D). Superficial layers in both dRSC and gRSC predominately contained PCs allocated to C3; which were cells with a high R_i_, a small hyperpolarisation-activated cation current (sag), and the fastest AP maximal rise time. This is consistent with superficial, late-spiking PCs with a relatively small soma reported by others [18]. In the deeper layers of the gRSC, most neurons belonged within C1, and therefore had the largest sag, greatest AP magnitude and lowest initial bursting of firing. This distribution of PC subtype is similar to those found in other murine cortical regions such as the prefrontal [26] and somatosensory cortices [27]. Interestingly however, cells in the dRSC deep layers were relatively evenly distributed between the cluster groups. In contrast to findings indicating PC homogeneity in some cortical areas [28], our findings suggest RSC contains diverse PC subtypes, which may confer distinct functional properties [18].

## Discussion

Here we have reported the relative selectivity of ACC, ATN and dSub inputs to both granular and dysgranular RSC. As far as we are aware, this is the first report looking at afferent inputs to dRSC at the synaptic level. The most striking observation was that all recorded pyramidal cells, both deep and superficial, in both RSC subdivisions, received a strong monosynaptic input from the ATN. ACC had lower connection probability than ATN and dSub and preferentially avoided superficial PCs in gRSC, which aligns with the one other published study describing RSC afferent inputs [22]. Although dSub axons were less dense in dRSC (Figure 1 and [22]), synaptic responses evoked by dSub axon activation were significantly *larger* in dRSC than gRSC. This mismatch between anatomical and functional connectivity is remarkable, and raises important questions about using anatomical tracing alone to infer connectivity. No evidence was found of pathway-specific differences in NMDA/AMPA ratios or overall PPR, although PPR did differ between sub-region and layer for dSub and AD/AV input. Therefore, the short-term plasticity of these projections differs dependent on the responding postsynaptic cell.

### Structural connectivity differences between presynaptic region inputs

The rodent RSC neuroanatomical connectivity has been well mapped, and some sub-region specific and laminar differences have been described [9–12,21,22]. However, tracing experiments in the present study quantified and described the pathway-specific laminar distribution of afferent terminals in both the gRSC and dRSC. Additionally, little work has been done to link anatomical and functional connectivity. Anatomically, ACC input preferentially targeted the dRSC over the gRSC, and fibres were most abundant in layer 6 for both sub-regions. Conversely, dSub and ATN input both preferentially targeted the gRSC and showed differences in laminar distribution between the sub-regions. In dRSC, dSub afferent terminals were densest in layer 6, whereas in gRSC, they primarily targeted layer 3. A similar pattern was found for the ATN afferent terminals; dRSC and gRSC layer 6 had the highest proportion of terminals but there was also an increase in targeting for layer 3 in the gRSC. The lamination is similar to that described in the rat brain [9–11], with some differences such as the layer 6 targeting by dSub and ATN. Additionally, little or no subicular input was found in the rat dRSC, whereas some fibres were observed in this study in the mouse.

The sub-region targeting preferences for each input remained consistent in the anterior and posterior RSC, however differences in overall fluorescent intensity along the anteroposterior axis were not measured as sections could not be normalised against each other. Certain pathways may preferentially target anterior or posterior regions of the RSC. Afferent and efferent connections with hippocampal structures vary depending on the RSC’s A-P position, as well as along the A-P and D-V axes in regions like the subiculum and CA1 [10]. Thalamocortical projections to the RSC from the ATN also preferentially target the anterior and posterior RSC depending on the location of the originating neurons along the A-P and D-V axes [29]. Furthermore, different information is relayed from the ACC, dSub and ATN to the RSC: for example, head direction signals are transmitted from the ATN [30] while the dSub conveys information about speed, place and trajectory [31].

In comparison, projections from the ACC to other cortical regions, such as the visual cortex, are responsible for top-down attentional control [32]. There is evidence that the anterior and posterior areas of the RSC are responsible for different aspects of spatial behaviour: the anterior RSC is suggested to be important for internally-directed navigation, while the posterior RSC is responsible for visually-guided navigation (reviewed by [3]). Moreover, the electrophysiological literature suggests that only neurons in the anterior RSC respond to location, direction and movement stimuli [3]. Therefore, it is probable that projection pathways conveying dissimilar forms of information will target the A-P axis differently. Our study has focused on afferent inputs to RSC from ‘higher’ cognitive regions involved in spatial memory and navigation, but it is important to note that RSC also receives inputs from multiple brain regions including primary sensory and motor cortices, and has been implicated in non-spatial memory formation (reviewed by [33]). Future research should also examine the role of sensory inputs in modulating RSC activity.

### Functional connectivity differences between presynaptic region inputs

The functional connectivity of the afferent inputs from the ACC, dSub and ATN differed in a variety of measures. Firstly, inputs from each presynaptic region had a differing probability of forming an excitatory synaptic connection with PCs in the RSC. Strikingly, stimulation of afferent terminals from the ATN resulted in EPSCs in every PC recorded, but inputs from the dSub and ACC were more sparsely distributed in the RSC. Approximately three-quarters of cells responded to dSub terminal stimulation, while fewer responded to ACC input, which formed connections with just over half of the recorded cells—still indicating a high degree of connectivity. This specificity of synaptic connection onto RSC PCs was modulated by layer and sub-region for ACC input, with most non-responders being found in the gRSC superficial layer, but not for dSub input. This specificity of synaptic connectivity, however, relates only to excitatory synaptic events as only EPSCs were recorded. While excitatory transmission is often the focus of long-range connectivity research, long-range inhibitory neurons have long been known to exist in cortical and hippocampal regions (reviewed by [34]), and a long-range inhibitory projection from CA1 to the RSC has recently been described [21].

The strength of the synaptic connectivity also differed between inputs: inputs from the ATN generated a median EPSC magnitude around 8-fold larger than those from the dSub and ACC. Therefore, the ATN pathway exerts a larger AMPA receptor-mediated excitatory effect on PCs in the RSC, but it is unknown whether this is due to more synaptic connections and/or an increase in postsynaptic AMPA receptors. A strong feed-forward drive of information from the ATN to RSC is important for head direction signalling [35], and a projection pathway with a strong input as seen is the present study would allow for transmission of this information with high fidelity. dRSC pyramidal cells had a lower probability of receiving input from dSub compared with gRSC, yet those neurons that did receive in input displayed a significantly larger magnitude of EPSC. Earlier research has shown that both dRSC and gRSC contribute to spatial memory but suggested that gRSC contributes to navigation using both internal and external cues, while dRSC is selectively involved in navigation when integrating spatial information [36]. The larger dSub EPSC in dRSC pyramidal cells may thus provide a wider temporal window to allow integration of spatial information with long-range inputs from different brain regions.

The NMDA receptor is a crucial component of excitatory synaptic transmission and long-term plasticity in the brain [37]. Its activation allows for Ca^2+^ influx into the neuron facilitating slower excitatory currents as well as regulating the synapse through various signalling pathways [38]. While ATN had the largest magnitude of NMDA-receptor mediated EPSCs, we found no difference in the ratio of NMDAR- to AMPAR-mediated responses between inputs, suggesting there is no significant difference in the proportion of NMDAR to AMPAR at the synapses for the different pathways. Studies examining pathways known to be highly plastic, such as CA3-CA1, normally observe NMDA/AMPA ratios between 0.6 and 1 (e.g. [39,40]). However, a much lower NMDA/AMPA receptor ratio was observed in all RSC afferent pathways examined, which corroborates previous findings that LTP cannot be induced in the RSC following tetanic stimulation [4]. Given that NMDA receptors in RSC are necessary for remote retrieval of context memory [41], the relatively low NMDA/AMPA ratio observed in our study raises interesting questions. Perhaps NMDA receptors play a more important role in intrinsic activity of RSC, rather than on long-range inputs?

Similarly, PPR was analysed as a measure of short-term plasticity; again, inputs from the different presynaptic regions did not differ significantly. On average, all inputs displayed PPD – signified by ratios of < 1 – and showed similar rates of recovery for EPSC magnitude following increased time between stimulations. PPD can be mediated by a variety of mechanisms, but most models suggest presynaptic vesicle depletion and postsynaptic receptor desensitisation (reviewed by [42]). This suggests that all tested RSC input pathways had comparable levels of glutamate release and rate of repletion [43,44] and/or similar AMPAR desensitisation [45]. This finding was unexpected as previous interrogation of these pathways in the gRSC showed that only the ATN pathway was depressing and that the ACC and dSub pathways were weakly facilitating [22]. However, in this study, PPR was recorded from a particular subtype of PCs (termed ‘low rheobase’ PCs) in current clamp mode, whereas synaptic recordings in the present study were performed under voltage-clamp conditions; as the neurons in our recordings were unable to fire action potentials, it was not possible to determine whether any of them were indeed low rheobase. Also, short-term synaptic dynamics are known to be cell subtype-specific [46] or even axon terminal-specific within the same neuron [47]. Close inspection of our data (Figure 5B) reveals variable PPR responses, with the range of PPR extending > 1. Thus, some cells in our sample showed paired pulse facilitation, although the average response was one of depression.

Differences in synaptic responses were also observed between RSC sub-regions and layers for some of the afferent input pathways. No effect of sub-region or layer was found for ACC input synaptic responses: PCs in the superficial and deep layers of the dRSC and gRSC all responded to ACC presynaptic excitation in a similar manner. However, synaptic strength did differ by sub-region for dSub and ATN inputs: for both pathways, cells in the dRSC displayed stimulus-evoked EPSCs of a greater magnitude than those in the gRSC. Although synaptic strength did not differ for either input between cells in the superficial and deep cortical layers. Finally, while all pathways showed PPD throughout the RSC, cells in the gRSC had a higher PPR and therefore weaker PPD than cells in the dRSC for both the ATN and dSub pathways. This is likely related to the finding that cells in the gRSC had a smaller synaptic response, suggesting presynaptic terminals in the dRSC may have a higher glutamate release probability. Furthermore, deep layer cells also showed a weaker PPD than cells in the superficial layers, but superficial and deep cells did not differ in the strength of their synaptic connections, so the mechanism is unknown. It is theorised that extent of synaptic depression is important for determining the neural code between PCs in the cortex [48], therefore the differences in PPD observed between RSC sub-regions and layers suggests different contributions of firing rates and temporal coherence.

### Pyramidal cell heterogeneity

Whilst PCs are the most common type of neuron in the mammalian neocortex, they exhibit a large diversity of subtypes with specific genetic profiles [49], circuitry [26], and morphological and electrophysiological properties [50]. Evidence from the rodent prefrontal cortex has shown that certain intrinsic physiological properties differ dependent on the location and efferent projections of PCs in the area [26]. Cortical PCs are a diverse neuronal population that have generally been divided into two sub-categories characterised by their firing properties: regular spiking (RS) and intrinsic bursting (IB) cells [51]. While most experiments examining PC diversity have used rodent models, RS and IB cells are found in similar proportions in the primate and human cortex [52–54]. Findings presented in this study confirm a similar diversity of pyramidal neurons in the retrosplenial cortex: a non-biased hierarchical clustering identifies three clusters (C1-C3) with distinct intrinsic passive and active membrane properties. C1 and C3 cells exhibited characteristics similar to RS cells, whilst C2 cells presented with a greater propensity for burst firing. However, other properties identified as significantly different in the C2 cluster are not necessarily associated with IB cells, such as a shorter AP halfwidth, faster 20-80% rise time and increased rheobase [55]. The relative lack of a hyperpolarisation-activated cation current (sag) shown by C2 cells is similar to that found in PCs in the superficial layers of other cortices [56], which could explain the high percentage of C2 cells found in the superficial layers of the RSC.

While the clustering of intrinsic properties shows some similarities to previously described PC sub-types, sufficient dissimilarities occur which prevent confident assignment to these sub-types. Furthermore, the three clusters described in the model only explained 30% of the variance. While the current data set could not be partitioned into more clusters without sacrificing cluster distinction, increasing the number of cells in the model could reduce variability and allow for further sub-categories of PCs to emerge within the broader classifications. The work here extends our knowledge of PC heterogeneity in the RSC by recording from cells in both the gRSC and dRSC. Interestingly, while latency to first AP was an important property for classifying PCs in the rat gRSC superficial layers [17,19], this variable was not significant in our model, similar to [18]. However, during the analysis no cluster of PCs exhibiting low rheobase [18] emerged either. This may be due to the method of analysis – a hierarchical clustering with no grouping pre-identified – requiring a larger sample size to reliably separate these cells.

### Concluding remarks

Here, we have examined the synaptic properties of afferent inputs to retrosplenial cortex from anterior cingulate cortex, dorsal subiculum and the anterior thalamic nucleus. Anterior thalamic connectivity appear ubiquitous, with every neuron tested displaying a post-synaptic response, irrespective of RSC subdivision or laminar location, that was often of large magnitude. We observed some surprising mismatches between anatomical and functional connectivity: gRSC received a far denser innervation of fibres from subiculum, but activation of these projections provided a stronger response in dRSC. Anatomical studies showing dense projections from dSub to gRSC [10,57] and those looking at Fos expression after spatial learning [e.g. 58] suggest that gRSC is the principal subdivision processing spatial information, although observations such as the development of place-cell like features in RSC [59] or the presence of spatial engrams in dRSC [60] challenge this view. Our data, showing a sparse but strong drive of dSub onto dRSC neurons, provides a circuits-based explanation for these observations. Whether or not GABAergic neurons in RSC display a similar unusual heterogeneity as pyramidal cells in this region, or whether they receive similar drive from retrosplenial afferents, remains to be seen.

For all the inputs studied, we found a relatively low contribution of NMDA receptors to the afferent-evoked EPSC. Future work studying the synaptic properties of intrinsic connections within RSC may explain why no researchers have yet reported the induction of LTP in a region associated with spatial memory. This could be a result of reduced NMDA receptors or perhaps a feature of inhibitory interneuron function in this area. Or it may be that previous studies have attempted to induce LTP via tetanic stimulation [4] when a form of spike-timing dependent plasticity may be required in RSC, perhaps via concurrent activation of the ubiquitous ATN projection alongside another pathway. Logically, one would expect a brain region implied in associational memory to be able to express LTP, but an STDP-like mechanism may well be more likely in a brain region receiving so many distinct inputs. The retrosplenial cortex remains an enigmatic brain region, presenting exciting challenges for future research on linking synaptic and cellular connectivity to behaviour.

## Methods

### Animals

All experiments were conducted in accordance with the UK Animals (Scientific Procedures) Act 1986 after local ethical review by the Animal Welfare and approved by the University of Exeter Institutional Ethical Review Board. We used both male and female C57BL/6J mice bred in-house at the University of Exeter for all experiments. All animals were maintained on a 12h constant light/dark cycle and procedures and experiments were conducted during the light phase. All animals had access to food and water *ad libitum* and were group-housed wherever possible. We used standard home-cage enrichment that included cardboard tubes, wooden chew blocks and nesting material.

### Drugs and Chemicals

CGP55845, Gabazine (SR 95531) and DNQX were purchased from HelloBio (UK), and all other chemicals were purchased from Sigma-Aldrich (UK) unless otherwise stated.

### Stereotaxic injection surgical procedure

All surgeries were conducted using aseptic technique. Mice were anaesthetised with isoflurane (5% induction, 1.2-2.5% maintenance) delivered in a constant flow of oxygen. The mice were placed on a heated pad (37°C) for the duration of the surgery and given 0.03 mg/kg of buprenorphine (buprenorphine hydrochloride, Henry Schein) subcutaneously at the beginning of surgery as an adjunct analgesic, plus 5 mg/kg of carprofen (Rimadyl, Henry Schein) and 10 ml/kg of 0.9% saline were given subcutaneously immediately post-surgery. 5 mg/kg of carprofen was also provided daily for 3 days post-surgery.

Mice received a unilateral 250 nl injection of viruses (at 100 nl/min; see below for vector details) in the ACC (AP: +1.75 mm; ML: ±0.20 mm; DV: −1.5 mm from pia), ATN (AP: −0.71 mm; ML: ±0.80 mm; DV −2.76 mm from pia) and/or dSub (AP: −3.79 mm; ML: ±2.25 mm; DV: −1.80 mm from pia). All co-ordinates are relative to Bregma in the head-flat position. After the surgery, the mice were allowed at least a 4-week recovery period to allow sufficient time for the expression of the viral construct.

### Anatomical tracing

We used C57BL6/J mice of both sexes aged between 3-5 months (mean age 3.5 months) in this tracing study. Mice were injected with either AAV8-hSyn1-chI-EGFP-WPRE-SV40p(A) (ETH Zurich Viral Vector Facility, titre 5.9 x 10^12^ vg/ml) or AAV8-hSyn1-chI-mCherry-WPRE- SV40(A) (ETH Zurich Viral Vector Facility, titre 5.6 x 10^12^ vg/ml) in two presynaptic brain regions (ACC, ATN and/or dSub). 6 weeks post-surgery, the mice were killed by transcardial perfusion / fixation with 4% paraformaldehyde (PFA) in 0.1 M phosphate buffered saline (PBS). Following perfusion, the brains were dissected out and post-fixed for 22 hours in 4% PFA solution, after which they were cryoprotected in 30% sucrose in 0.1 M PBS. 50-micron coronal sections were taken using a freezing sledge microtome (Leica SM2010R with Physitemp BFS-5MP temperature controller). Selected slices containing the anterior RSC, posterior RSC, ACC, ADAV and dSub were mounted using Hard Set mounting medium with DAPI (Vectashield, 2BScientific). Fluorescent fibres were visualised using epifluorescent microscopy (Nikon Eclipse 800 microscope with CoolLED pE-4000 LED light source), and images were captured using a SPOT RT monochrome camera running SPOT Basic imaging capture software (SPOT imaging) at 10x magnification. Duplicate images of the anterior and posterior RSC were taken for each injection. Mean fluorescence intensities as compared to total image fluorescence were calculated for the gRSC and dRSC using Fiji software [61], and then for each cortical layer, in each RSC section and averaged. As mean fluorescent intensities were used for analysis, and both eGFP and mCherry fluorophores were used equally across sources of interest, fluorophore was not controlled for.

18 mice (n_male_ = 8, n_female_ = 10) were initially injected, and 7 mice were excluded completely from analysis due to failed or misplaced injections in both target regions. Of the 11 remaining mice (n_male_ = 4, n_female_ = 7); 7 mice had one correctly placed injection and 4 mice had two correctly placed injections leaving 15 positive injection sites of which 6 were in the ACC, 4 in the dSub and 5 in the AD/AV.

### Slice preparation and electrophysiology

For the optogenetic synaptic recordings, we used mice of both sexes aged between 4-8 months (mean age 6.8 months) at time of recording. Mice were unilaterally injected with AAV_5_-hSyn1-ChR2(H134R)-mCherry-WPRE-hGHp(A) (ETH Zurich Viral Vector Facility, titre 7.1 x 10^12^ vg/ml) 14 mice (n_male_ = 7, n_female_ = 7) had correctly placed injections in the ACC; from which 123 cells were recorded and 30 then excluded from analysis (see data analysis). 10 mice (n_male_ = 6, n_female_ = 4) had correctly placed injections in the dSub; from which 84 cells were recorded and 19 excluded. Finally, 9 mice (n_male_ = 7, n_female_ = 2) had correctly placed injections in the ADAV; from which 52 cells were recorded and 13 excluded. In addition, 2 mice were injected in the ADAV with the control virus AAV_8/2_-hSyn1-mCherry used in the anatomical tracing experiments; from which 21 cells were recorded and 1 cell excluded.

For the neuronal intrinsic properties recordings we used 27 mice (n_male_ = 12, n_female_ = 15) aged between 3-6 months (mean age 5.7 months): from which 163 cells were recorded, and 35 were excluded.

Mice were anaesthetised with isoflurane (5%) and decapitated before the brain was rapidly dissected out and placed in room temperature oxygenated (with 95% O_2_, 5% CO_2_ carbogen gas, BOC gases) NMDG cutting solution, containing (in mM): 135 NMDG, 10 D-Glucose, 1.5 MgCl_2_, 0.5 CaCl_2_, 1 KCl, 1.2 KH_2_PO_4_, 20 Choline bicarbonate. RSC-containing 300 µm coronal slices corresponding to approximately Bregma −0.5 mm to −4.5 mm were sectioned using a Leica VT1200 vibratome. Following sectioning, slices were transferred to a holding chamber containing artificial cerebral spinal fluid (aCSF) perfused with a continuous flow of carbogen (95% O_2_, 5% CO_2_), and containing (in mM): 119 NaCl, 3 KCl, 1 NaH_2_PO_4_, 26 NaHCO_3_, 10 D-Glucose, 2.5 CaCl_2_, 1.3 MgCl (pH 7.4). The slices were incubated in the holding chamber at 35°C for 30 minutes, then at RT for at least another 30 minutes before recording. For optogenetic experiments, slices containing the injection site were also collected and injection site fluorescence was visually confirmed immediately following slicing.

RSC containing slices were attached to a 0.1% poly-L-lysine coated coverslip and transferred to the recording chamber where they were perfused with carbogen-saturated aCSF (4 ml/min, 32-34°C). Differential interference contrast and fluorescent signal imaging was perfomed using an Olympus BX51W1 microscope and SciCam Pro camera with a CoolLED light source. Recording microelectrodes were fabricated from borosilicate glass (World Precision Instruments; OD 1.5 mm, ID 0.86 mm, 3-6 mΩ) and filled with solution. PNs were tentatively identified under differential interference contrast visualisation, and assigned as being located in the superficial (supragranular) or deep (infragranular) layers of the dRSC or gRSC. Confirmation of cell-type and location was conducted for a sub-set of cells by visualising cell morphology via *post-hoc* cell recovery. All electrophysiological recordings were acquired using an Axon Multiclamp 700B (Molecular Devices, USA), digitally sampled at 20 kHz with a low-pass filter of 8 kHz on an Axon Digidata 1440A (Molecular Devices, USA) and sampled on a PC running pClamp 9 (Molecular Devices). The liquid junction potential was not corrected.

Intrinsic properties experiments were recorded in current clamp mode using a K-gluconate intracellular solution, containing and (in mM): 135 K-gluconate, 3 MgCl_2_, 0.5 EGTA, 10 HEPES, 0.3 Na_2_-GTP, 2 Mg-ATP plus 2 mg/ml biocytin. Cells were recorded at V_m_ with no injection of bias current, and all recordings were made in standard aCSF. Rheobase was measured using a 200 ms square-wave positive current injection protocol incrementing in 2 pA steps until an AP was induced. All other measures were computed from a 1000 ms square-wave negative and positive current injection protocol incrementing in 50 pA steps.

Synaptic properties experiments were recorded in voltage clamp mode using a caesium methanesulphonate (CsMeSO_4_) intracellular solution containing (in mM): 135 CsMeSO_4_, 8 KCl, 0.5 EGTA, 10 HEPES, 0.5 QX-314, 0.1 Spermine, 0.3 Na_2_-GTP, 2 Mg-ATP. Cells were recorded from at a holding membrane potential (V_H_) of −70 mV for recordings unless otherwise stated. All recordings were made in standard aCSF unless otherwise stated. All optic stimulation was carried out at a 470 nm wavelength generated by a CoolLED pe-4000 light source at 100% power.

Cell response and EPSC magnitude were measured using a stimulation train protocol consisting of a 500 ms −10 mV square voltage injection followed by 5 ms optic stimulation pulses at 7 Hz frequency applied to the cell for 10-20 sweeps. PPR was measured using an increasing inter-pulse interval (IPI) protocol which consisted of: a 500 ms −10 mV square voltage injection followed by two 5 ms optic stimulation pulses at 10, 17, 51, 100, 170, 510 and 1000 ms intervals. Finally, AMPAR- and NMDAR-mediated currents were measured using a protocol consisting of a 500 ms −10 mV square voltage injection followed by a single 5 ms optic stimulation pulse. AMPAR-mediated currents were measured at V_H_ = −70 mV in standard aCSF and NMDAR-mediated currents were measured at V_H_ = +40 mV in aCSF containing antagonists for GABA_A_, GABA_B_ and non-NMDA ionotropic glutamate receptors (standard aCSF containing (in µM): 10 Gabazine, 1 CGP-55845, 10 DNQX).

### Data Analysis

From the current clamp recordings, the following intrinsic neuronal properties were calculated: resting membrane potential (V_m_), input resistance (R_i_), membrane time constant (τ), hyperpolarisation-activated cation current-mediated sag and rebound, spiking threshold, latency to first AP, AP amplitude, AP halfwidth, AP maximal rate of rise, accommodation index (AI) and burst index (BI). Cells were excluded from analysis if they had an access resistance > 20 MΩ or V_m_ > −50 mV.

Resting membrane potential (V_m_) was calculated as the average membrane voltage for 100 ms before current injection. Input resistance (R_i_) was calculated as mean voltage deflection during the last 200 ms – the steady state response (ssV) – of the first hyperpolarisation current injection sweep divided by the amplitude of the negative current injection. Membrane time constant (τ) was calculated using a first order exponential curve fitted to the charge curve of the membrane between 20 and 80% of the peak amplitude of voltage deflection. Sag and rebound voltages were computed from the first hyperpolarisation injection. Sag was calculated as percentage of the difference between the maximum negative voltage deflection (V_min_) and steady state response (ssV) divided by the difference between V_min_ and V_m_. Rebound voltage was calculated as the difference between the maximum positive voltage peak following the end of the negative current injection step (V_max_) and V_m_.

Spike characteristics were computed from the first AP of the first spike train to contain ≥4 spikes. AP threshold was calculated as the voltage at which the first derivative of the membrane potential (dV/dt) exceeded 20 mV/ms. Maximal rate of rise (max dV/dt) was calculated as the peak of the first derivative of the membrane potential. AP peak was computed as the absolute maximum voltage of the AP waveform. AP half-width was calculated as the width at half maximal voltage from AP threshold. Burst index (BI) was calculated as the sum of the inverse of the spike number squared multiplied by the ISI. Rheobase was calculated by manually identifying the depolarising current injection magnitude when an AP was first generated on ClampFit software (Molecular Devices).

From the voltage clamp recordings the following synaptic properties were calculated: binary response rate, EPSC magnitude, PPR and NMDA/AMPA ratio. Cells were excluded from analysis if they had an access resistance of > 20 MΩ or displayed a change in R_i_ (ΔR_i_) > 20%. R_i_ was calculated for each sweep as the mean current deflection during the voltage injection step divided by the amplitude of the negative voltage injection. ΔR_i_ percentage was computed by subtracting the R_i_ of the first sweep from the final sweep, then dividing by first sweep R_i_. Average R_i_ was calculated as the mean R_i_ for all sweeps.

EPSC magnitude was calculated from the first optogenetic stimulation pulse generated in the 7 Hz stimulation protocol. Cells were excluded if <10 viable sweeps were recorded. Sweeps were averaged and normalised: baseline current was calculated as the mean current for the 500 ms following voltage injection, and adjusted to 0 pA. The binary response rate threshold for monosynaptic cell connectivity was set at > 2 pA positive or negative inflection and onset within the 5 ms optogenetic stimulation window. The EPSC magnitude was calculated as either the maximum negative current peak or the current at time of the first EPSC peak; where a multi-peak response generated defined peaks before the maximum. Identification of peaks before maximum was computed by plotting the first order differential of the response and identifying peaks of greater than 4 standard deviations from the differential trace.

Three replicate recordings were made for each cell for the PPR protocol, and corresponding sweeps were averaged together. R_i_ calculation and baseline adjustment were computed as described above. EPSC magnitude for each pulse was calculated as the maximum negative current peak, and a PPR ratio was generated for each averaged sweep by dividing the second pulse EPSC by the first pulse EPSC. Sweeps used to compute AMPAR- and NMDAR-mediated currents were averaged and normalised for each block. R_i_ and ΔR_i_ were calculated for the AMPAR-current sweeps only.

AMPAR-mediated EPSC magnitude was calculated as the maximum negative current peak in a 200 ms window following optogenetic stimulation. NMDAR-mediated EPSC magnitude was calculated as the maximum positive current peak in the same time window. AMPA-NMDA ratio was then calculated by dividing NMDAR EPSC by the AMPAR EPSC. Any cells which displayed AMPAR, but not NMDAR, responses were discarded from analysis.

Electrophysiological data analysis was carried out using custom MATLAB (Mathworks) scripts unless stated otherwise. Representative electrophysiological traces were generated using Igor Pro 7 (Wavemetrics).

### Statistical Analysis

RStudio [62] was used to conduct all statistical analyses and generate all data graphs. All data was tested for normality where necessary, and parametric or non-parametric tests were used where appropriate. When comparing between groups independent and mixed ANOVA were used or Kruskal-Wallis and Scheirer-Ray-Hare tests, which were followed by *post hoc* Mann-Whitney U or t-test multiple comparisons respectively. Synaptic response probability was analysed using Pearson’s χ^2^ test, followed by *post hoc* χ^2^ tests.

Mixed effect models were used to compare synaptic response data between RSC sub-regions and layers. These models included mouse ID as a random effect (random slope and intercept) to control for multiple cells being recorded from the same animal within a presynaptic input group. For EPSC magnitude and AMPA/NMDA ratio, a null model was fitted with just mouse random effect, then a linear mixed effect model was fitted by force-entering all independent variables as fixed effects. For PPR data, the null model included mouse random effect and IPI ratios as a repeated fixed effect factor. The goodness-of-fit for the null and experimental models was compared using the χ^2^-likelihood ratio test, and the mixed model was reported when it generated a significant improved upon the null model. Goodness-of-fit was also assessed and reported using the Akaike information criterion (AIC). Main effects were calculated using ANOVA, and pairwise comparisons reported as the fixed effect estimate and t-test statistics. Where random effects were found, the contribution to variance was reported.

For the intrinsic membrane properties data, the variables were pre-processed by scaling between 0 and 1, before hierarchically clustering using Euclidean distances and Ward’s method. The hierarchical cluster was then cut: cluster n was chosen by considering multiple cluster index measures (such as the Hubert and silhouette indices). Hierarchical clustering bootstrapping (1000 runs) using the Jaccard coefficient [63] was then used to test cluster stability. Pearson’s r correlation coefficient was used to test for collinearity on the non-standardised variables. Due to multiple significant correlations, a multifactorial ANOVA was conducted to examine difference between the clusters, followed by *post hoc* ANOVA and Tukey’s HSD tests.

All *post hoc* analyses were *p*-adjusted using the Benjamini-Hochberg correction to control for false discovery rate, unless specified as Tukey’s HSD tests. Mice of both sexes were used in all experiments, but sex was not controlled for.

## Acknowledgements

This work was supported by Biotechnology and Biological Sciences Research Council grant BB/P001475/1 (MTC). Salary support from SK was provided by Alzheimer’s Research UK Interdisciplinary research grant ARUK-IRG2017B-4 (MTC). GMS was a GW4 BioMed doctoral training program student funded by the Medical Research Council (MR/N0137941/1). For the purpose of open access, the authors have applied a Creative Commons Attribution (CC-BY) licence to any Author Accepted Manuscript version arising from this submission. We are grateful to Dr Michael M Kohl (University of Glasgow) for discussions on the physiology and function of retrosplenial cortex.

## Author Contributions

Conceptualization, G.M. and M.T.C.; Methodology, G.M. and M.T.C.; Investigation, G.M., L.A., and S.K.; Writing – Original Draft, G.M., and M.T.C.; Funding Acquisition, M.T.C., J.P.A.; Resources, M.T.C. and A.R.; Supervision, M.T.C., A.R., J.P.A., and J.W.

