## Supplementary material for "Dissection of retrosplenial cortex inputs: ubiquitous drive from anterior thalamus"

Supplementary Figures

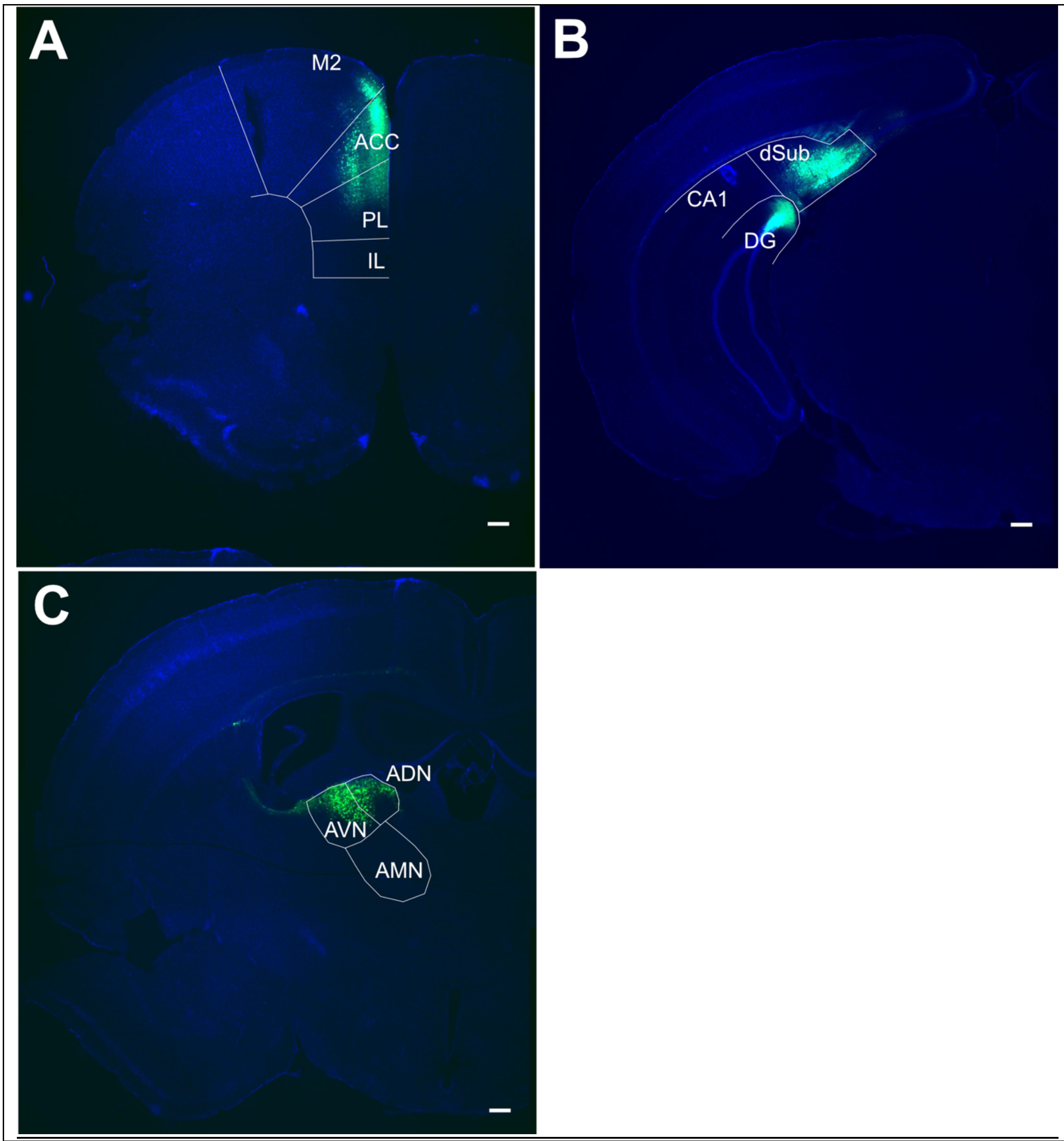


**Supplementary figure 1: Representative images of injection sites from anatomical tracing experiments. A,** injection site in anterior cingulate cortex (ACC). **B**, injection site in dorsal subiculum. **C**, injection site in AD/AV thalamus. Scale bar: 250 µm

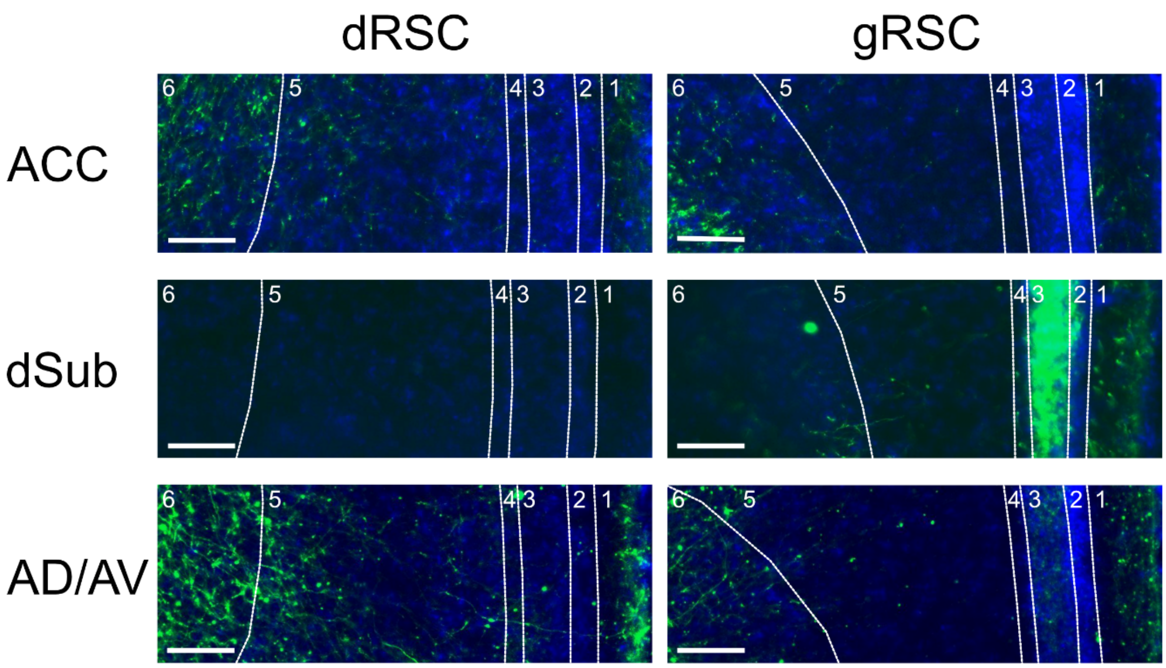


**Supplementary figure 2: Magnified representative images of afferent terminals from anatomical tracing experiments.** Cross-section of anterior gRSC and dRSC images. Scale bar: 100 µm

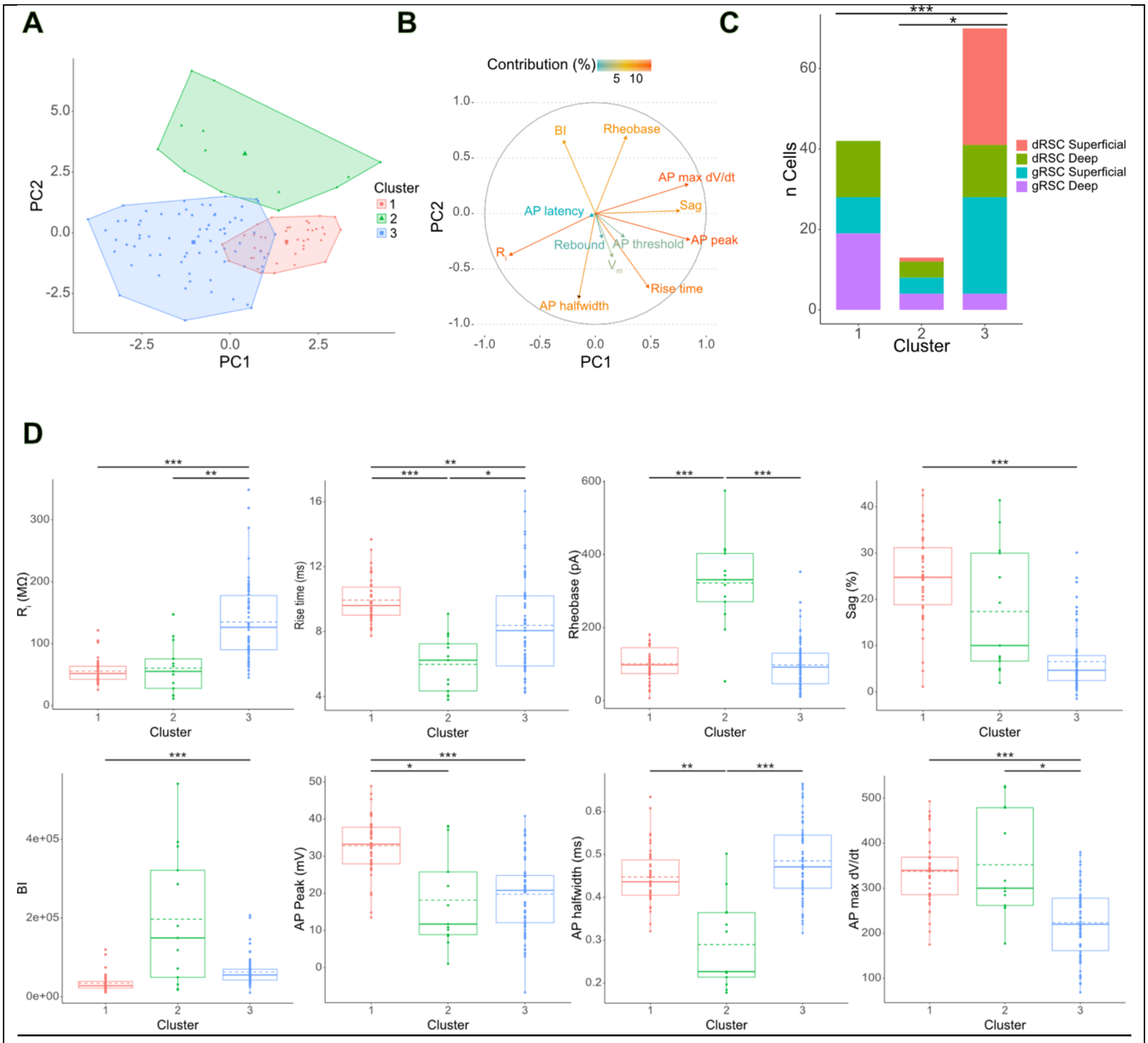


**Supplementary figure 3: further cell clustering analysis. A,** scatter plot of principal components PC1 and PC2 with hierarchical clusters imposed. **B,** variable loadings onto PC1 and PC2. **C**, There were significantly 76 different proportions of PCs from the different sub-regions of the RSC between the clusters ($\chi 2(6) = 43.92$, $p <$ .001; Pearson's Chi-Squared test). Chi-squared *post hoc* tests showed that the cell proportions between C1 and C2 did not differ significantly ($p$ =0.25), but did differ between C1 and C3 and C2 and C3. * *p < .05, ** *** *p < .001*. **D,** cell clusters have distinct differences in intrinsic membrane properties. Significant differences between clusters were seen for $R_i$, rise time, rheobase, sag, BI, and AP peak, half-width and max dV/dt. Supplemental Table 3 provides a breakdown of statistics and significances. Boxplots display median (solid line), mean (dashed line), IQR and range. * *p < .05, ** p < .01, *** p < .001*

Supplementary Tables

| | EPSC magnitude | | | | | | | | |
|---|---|---|---|---|---|---|---|---|---|
| | **ACC** | | | **dSub** | | | **ADAV** | | |
| *Predictors* | *Estimates* | *t* | *p* | *Estimates* | *t* | *p* | *Estimates* | *t* | *p* |
| Intercept | 112.71 (64.84 – 160.57) | 4.78 | **<0.001** | 211.07 (115.03 – 307.11) | 4.44 | **<0.001** | 684.53 (467.91 – 901.15) | 6.42 | **<0.001** |
| Sub-region: gRSC | -34.05 (-94.42 – 26.31) | -1.14 | 0.26 | -132.55 (-233.79 – -31.31) | -2.64 | **0.012** | -351.81 (-606.12 – -97.51) | -2.81 | **0.008** |
| Layer: Deep | -35.03 (-96.31 – 26.25) | -1.16 | 0.254 | 5.99 (-79.46 – 91.45) | 0.14 | 0.888 | -111.06 (-367.43 – 145.30) | -0.88 | 0.385 |
| **Random Effects** | | | | | | | | | |
| $\sigma^2$ | 6543.56 | | | 17196.91 | | | 140021.34 | | |
| $\tau^2_{mouse}$ | 1358.64 | | | 0 | | | 14039.34 | | |
| ICC | 0.17 | | | | | | 0.09 | | |
| N | 13 | | | 9 | | | 9 | | |
| Observations | 41 | | | 47 | | | 39 | | |
| *Marginal $R^2$ / Conditional $R^2$* | 0.096 / 0.251 | | | 0.136 / NA | | | 0.199 / 0.272 | | |

**Supplementary Table 1. Mixed model results for EPSC magnitude**

The mixed effect model did not significantly improve upon the null model for ACC input ($\chi_2(2)$ = 3.7, *p* = .16; **AIC$_{null}$** = 492.4, **AIC$_{mem}$** = 492.7), but did improve for dSub ($\chi_2(2)$ = 6.7, *p* < .05; **AIC$_{null}$** = 604.5, **AIC$_{mem}$** = 601.7) and ADAV input (($\chi_2(2)$ = 8.2, *p* < .05; **AIC$_{null}$** = 590.2, **AIC$_{mem}$** = 586.0).

For fixed effects: table displays effect sizes, confidence intervals, *t* statistic and significant values. For random effects: table displays residual variance ($\sigma^2$), mouse variance ($\tau^2$), and the intraclass correlation coefficient (ICC). Marginal $R^2$ refers to variance explained by fixed effects only, and conditional $R^2$ to variance explained by combined fixed and random effects.

| | PPR | | | | | | | | |
|---|---|---|---|---|---|---|---|---|---|
| | **ACC** | | | **dSub** | | | **ADAV** | | |
| *Predictors* | *Estimates* | *t* | *p* | *Estimates* | *t* | *p* | *Estimates* | *t* | *p* |
| Intercept | 0.44 (0.21 – 0.67) | 3.76 | **<0.001** | 0.02 (-0.15 – 0.19) | 0.23 | 0.819 | 0.21 (0.05 – 0.37) | 2.66 | **0.009** |
| Sub-region: gRSC | 0.06 (-0.08 – 0.20) | 0.82 | 0.412 | 0.27 (0.18 – 0.37) | 5.56 | **<0.001** | 0.14 (0.04 – 0.25) | 2.67 | **0.008** |
| Layer: Deep | 0 (-0.16 – 0.16) | -0.03 | 0.976 | 0.19 (0.09 – 0.30) | 3.77 | **<0.001** | 0.18 (0.08 – 0.28) | 3.55 | **0.001** |
| PPR: 17 ms | -0.06 (-0.30 – 0.18) | -0.52 | 0.604 | 0.06 (-0.09 – 0.20) | 0.74 | 0.461 | 0.08 (-0.10 – 0.26) | 0.92 | 0.361 |
| PPR: 51 ms | 0.21 (-0.03 – 0.45) | 1.72 | 0.087 | 0.33 (0.18 – 0.47) | 4.37 | **<0.001** | 0.29 (0.11 – 0.47) | 3.18 | **0.002** |
| PPR: 100 ms | 0.28 (0.05 – 0.52) | 2.34 | **0.02** | 0.27 (0.12 – 0.41) | 3.58 | **0.001** | 0.24 (0.06 – 0.42) | 2.61 | **0.01** |
| PPR: 170 ms | 0.39 (0.15 – 0.63) | 3.2 | **0.002** | 0.22 (0.07 – 0.37) | 2.91 | **0.004** | 0.19 (0.01 – 0.37) | 2.08 | **0.039** |
| PPR: 510 ms | 0.3 (0.06 – 0.54) | 2.46 | **0.015** | 0.23 (0.08 – 0.37) | 3.04 | **0.003** | 0.23 (0.05 – 0.41) | 2.54 | **0.012** |
| PPR: 1000 ms | 0.43 (0.20 – 0.67) | 3.58 | **<0.001** | 0.42 (0.27 – 0.57) | 5.62 | **<0.001** | 0.42 (0.24 – 0.60) | 4.56 | **<0.001** |
| **Random Effects** | | | | | | | | | |
| $\sigma^2$ | 0.22 | | | 0.05 | | | 0.11 | | |
| $\tau^2_{mouse}$ | 0.02 | | | 0.01 | | | 0 | | |
| ICC | 0.09 | | | 0.22 | | | 0.01 | | |
| N | 8 | | | 6 | | | 8 | | |
| Observations | 210 | | | 126 | | | 182 | | |
| *Marginal $R^2$ / Conditional $R^2$* | 0.114 / 0.197 | | | 0.401 / 0.535 | | | 0.200 / 0.210 | | |

### Supplementary Table 2. Mixed model results for PPR

The mixed effect model did not significantly improve upon the null model for ACC input ($\chi_2(2) = 0.8$, $p = .69$; **AIC$_{null}$** = 299.1, **AIC$_{mem}$** = 302.3), but did improve for dSub ($\chi_2(2) = 35.2$, $p < .001$; **AIC$_{null}$** = 35.4, **AIC$_{mem}$** = 4.2) and ADAV input ($\chi_2(2) = 18.1$, $p < .001$; **AIC$_{null}$** = 141.9, **AIC$_{mem}$** = 127.8).

| | NMDA/AMPA Ratio | | | | | | | | |
|---|---|---|---|---|---|---|---|---|---|
| | **ACC** | | | **dSub** | | | **ADAV** | | |
| *Predictors* | *Estimates* | *t* | *p* | *Estimates* | *t* | *p* | *Estimates* | *t* | *p* |
| Intercept | 0.16<br>(-0.07 – 0.39) | 1.53 | 0.15 | 0.34<br>(0.03 – 0.64) | 2.82 | 0.037 | 0.29<br>(-0.07 – 0.66) | 1.88 | 0.102 |
| Sub-region: gRSC | 0.1<br>(-0.18 – 0.39) | 0.78 | 0.449 | -0.17<br>(-0.45 – 0.10) | -1.63 | 0.164 | 0.13<br>(-0.28 – 0.53) | 0.75 | 0.476 |
| Layer: Deep | 0.06<br>(-0.24 – 0.36) | 0.42 | 0.679 | 0.1<br>(-0.12 – 0.32) | 1.18 | 0.29 | 0.02<br>(-0.33 – 0.36) | 0.11 | 0.915 |
| **Random Effects** | | | | | | | | | |
| $\sigma^2$ | 0.07 | | | 0.01 | | | 0.05 | | |
| $\tau^2_{mouse}$ | 0 | | | 0 | | | 0.01 | | |
| ICC | | | | | | | 0.19 | | |
| N | 9 | | | 7 | | | 8 | | |
| Observations | 18 | | | 10 | | | 12 | | |
| *Marginal $R^2$ / Conditional $R^2$* | 0.068 / NA | | | 0.470 / NA | | | 0.053 / 0.238 | | |

**Supplementary Table 3. Mixed model results for NMDA/AMPA ratio**

The mixed effect model did not significantly improve upon the null model for neither ACC ($\chi_2(2) = 1.2$, $p = .55$; **AIC$_{null}$** = 9.2, **AIC$_{mem}$** = 12), dSub ($\chi_2(2) = 5.9$, $p = .05$; **AIC$_{null}$** = -2.6, **AIC$_{mem}$** = -4.5), nor ADAV input ($\chi_2(2) = 0.5$, $p < 0.78$; **AIC$_{null}$** = 7.3, **AIC$_{mem}$** = 10.8).
